# Learning with interacting dendrites improves neuronal familiarity detection

**DOI:** 10.64898/2026.08.20.746078

**Authors:** Fangxu Cai, Marcus K. Benna

## Abstract

Biological neurons can perform nonlinear computations within their dendrites and support branch-localized plasticity. This raises the possibility that single cells can store memories more efficiently and with less interference by confining synaptic modifications to specific dendrites. We study a parallel-dendrite model performing online familiarity detection and compare three dendrite-update rules during learning: (i) independent thresholding, (ii) an interacting rule that adapts the target local dendritic activation per item, and (iii) an interacting n-winners-take-all (WTA) rule that constrains the number of updated branches per item. The interacting rules substantially improve capacity by limiting variance in memory responses and decorrelating weights across branches — even when inputs are strongly correlated. These results suggest that competition among dendrites, consistent with resource-limited plasticity mechanisms, can enhance single-cell memory beyond non-interacting schemes.

## Introduction

Dendrites endow neurons with rich nonlinear processing capabilities, enabling local subunit computations that transform how inputs are integrated and learned [1, 2, 3, 4]. Prior experimental and modeling results suggest that dendrites can operate as semi-independent nonlinear units, with a neuron acting like a two-layer network whose first layer consists of dendritic subunits and whose second layer sums the subunit outputs at the soma [5, 6, 7, 8]. Neurons described by such models have enhanced computational power compared to a more standard point-neuron model that linearly integrates inputs (and then applies a simple nonlinear activation function at the soma, which may be present in both types of models).

The computational role of dendrites has been studied at multiple levels of abstraction [9]. At the simplest end, reduced two-layer and subunit models represent a neuron as a set of nonlinear dendritic units whose outputs are summed at the soma [5, 6]. More biophysically motivated but still simplified models show that passive dendritic cable properties can enable linearly non-separable computations [10], and that sublinear dendritic integration can similarly expand single-neuron computations [11]. Other models incorporate additional dendritic mechanisms, showing that dendritic nonlinearities can support efficient spike-based computations under in-vivo-like input statistics [12], that active dendrites can allow sparse strong inputs to determine stimulus selectivity [13], and that compartmentalized dendritic pooling can generate orientation selectivity [14]. More detailed biophysical models examine how dendritic morphology, synapse location, input timing, active conductances, and local ionic dynamics shape integration and computation [9]. Recent studies have used such models to explain dendritic action potentials and nonlinear computations in human cortical pyramidal neurons [15], cell-type-specific dendritic integration in retinal ganglion cells [16], and how local ion dynamics can regulate input integration in active dendrites [17]. Studies of dendritic learning further investigate how plasticity can exploit these dendritic computations. Dendritic prediction of somatic spiking can provide a local learning signal [18]; somato-dendritic plasticity can approximate error-backpropagation in active dendrites [19]; somatic- and dendritic-spike-mediated plasticity can be modeled together in single neurons and networks [20]; and cable-theory-derived synaptic learning rules can allow detailed pyramidal-neuron models to learn nonlinear dendritic computations [21]. Together, these studies show that different levels of dendritic detail can be used to study different aspects of neuronal computation, from nonlinear integration and stimulus selectivity to branch-local learning rules.

A wealth of experimental evidence, largely driven by in-vivo two-photon calcium imaging, has demonstrated branch-specific selectivity. In the motor cortex, for instance, learning distinct motor tasks drives activity and persistent synaptic plasticity on different, non-overlapping branches of the same pyramidal neuron [22]. In the visual cortex, individual basal dendrites of a single neuron can be tuned to different orientations of a visual stimulus [23]. In the hippocampus, recent work has provided evidence for branch-specific place tuning, where different dendritic branches of a CA3 neuron can encode distinct spatial locations [24]. By spatially segregating memories onto different branches, a single neuron can participate in multiple memory traces simultaneously while minimizing interference between them.

Complementing these experimental findings, theoretical and computational models have established that sparse, localized plasticity is a highly efficient strategy for information storage [25, 26]. Memory capacity can be greatly increased if learning events modify only a small fraction of a neuron’s total synapses, a condition naturally met by confining plasticity to one or a few dendritic branches. This principle raises a critical question: if only a subset of dendrites are engaged to store a new memory, what mechanism governs their selection? The simple two-layer dendritic memory models typically treat dendrites as independently thresholded learning units, so that branch recruitment is determined independently by local activity [27, 26]. Interactions or competition between dendritic branches can arise in more mechanistic models [28, 29], but such models often include multiple biophysical or plasticity mechanisms, making it harder to isolate the computational benefit of the branch-level interaction itself. Here, we use a minimal two-layer parallel dendrite model to study how explicit interactions among dendrites during branch selection affect memory storage. We propose and test two interacting learning rules: a variable- *u*_target_ rule that adaptively sets the per-item update target for each dendritic branch, and an “n-winners-take-all” selection rule, where dendrites actively compete and only a fixed number of the most highly activated branches are recruited for synaptic modification. We investigate the computational consequences of these interacting rules compared to a non-interacting, threshold-based rule within a parallel dendrite model performing an online familiarity detection task. This work thus helps elucidate how the strategic selection of plastic subunits can contribute to maximizing the storage capacity of individual neurons.

### Task and Model

To quantify the memory capacity of a single neuron, we employ an online familiarity detection task, following a similar approach to [26]. In this task, a neuron is sequentially presented with unique input patterns for one-shot learning (Fig. 1a, left). The neuron is expected to generate a high memory response for recently presented patterns (familiar) and a low response for patterns presented long ago (unfamiliar). Fig. 1a (right) illustrates a typical memory response distribution as a function of pattern age. The shaded blue region represents the distribution of the memory response, spanning from the 1st to the 99th percentile. Memory capacity is defined as the pattern age when the 1st percentile of the familiar response distribution intersects with the 99th percentile of the unfamiliar (steady-state) response distribution. This intersection point corresponds to the number of subsequently stored patterns at which the false positive and false negative error rates are both 1%.

**Figure 1.**
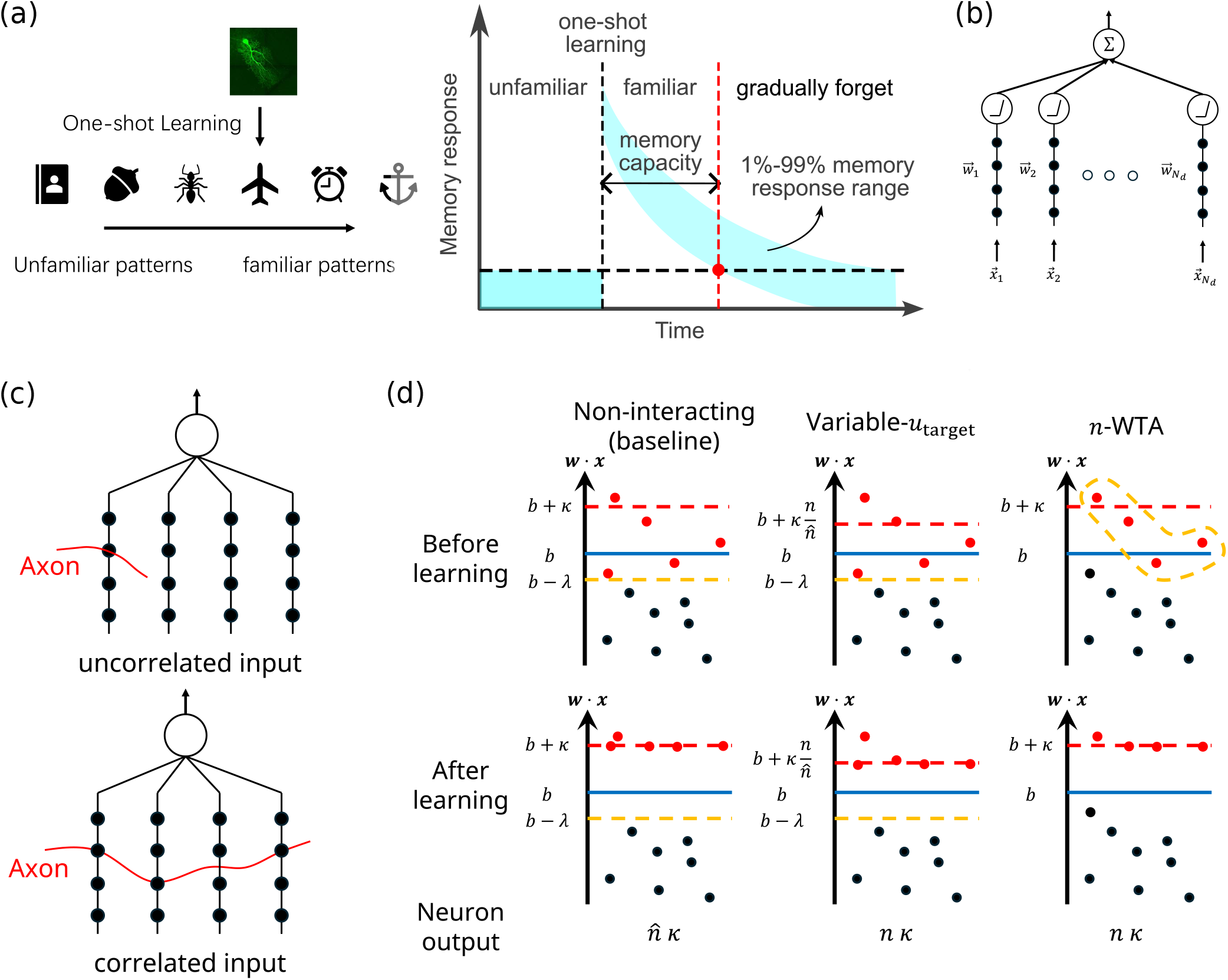
Task and model setup. (a) Left: Illustration of the online familiarity detection task. Input patterns are presented sequentially to the neuron, which performs one-shot learning on each of them. Right: Schematic of memory response distribution as a function of pattern age. Memory capacity is defined as the age when the 1% lower bound (1st percentile) of the transient response intersects with the 99% upper bound of the steady state response. (b) Schematic of the parallel dendrite model. (c) Two connectivity patterns. Top: an axon only makes one connection with the neuron; inputs are uncorrelated. Bottom: an axon can make multiple connections, but only one per dendrite; inputs are correlated across dendrites. (d) Three dendrite selection rules. Left: a non-interacting threshold-based rule. Middle: a variable-target rule in which the desired dendritic activation can change as needed to maintain a constant neuronal output. Right: an interacting n-winners-take-all (n-WTA) rule where the top n dendrites are selected. The selected dendrites undergo weight changes to achieve a target activation level.

The two-layer parallel dendrite model [5] we use is depicted in Fig.1b. In this model the detailed neuronal morphology is ignored, and *N*_*d*_ dendrites, each with *N*_*s*_ synapses, are connected in parallel, resulting in a total of *N*_tot_ = *N*_*d*_ *× N*_*s*_ synapses. For the *r*-th dendrite, we denote the synaptic weight vector as 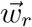 and its corresponding input vector as 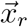. A complete input pattern is therefore represented by the set of vectors 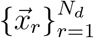. When an input pattern is presented, each dendrite computes a local activation, 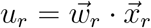. These local activations are then passed through a non-linear function (ReLU) and summed at the soma to generate the total memory response, *Y*:

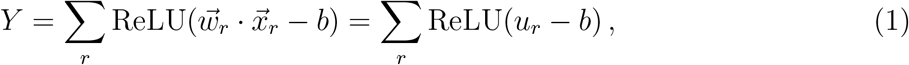

where *b* is the activation threshold of each dendrite. The resulting neural activity in general is a non-linear function of *Y*.

Input patterns received by the dendritic tree are modeled to possess a fixed correlation structure, which is defined by the connectivity between incoming axons and the postsynaptic dendrites. We assume that signals carried by individual input axons are uncorrelated, but a single axon can form multiple synapses with the dendritic tree, thereby inducing correlations in the inputs the dendrites receive. Specifically, we investigate two scenarios, as shown in Fig. 1c: no correlation (top), where each axon forms only one synapse with the entire dendritic tree, and cross-dendrite correlation (bottom), where an axon can form multiple synapses, but with at most one synapse per dendrite. To quantify the input correlation between dendrites, we define a shared-input coefficient *c* = *N*_*a*_*/N*_*s*_, where *N*_*a*_ is the number of axons available to a neuron (not all available axons necessarily have to make connections with the neuron). A smaller *c* value corresponds to a higher chance that two dendrites make connections with the same axon. The probability that, for a pair of dendrites, an axon that forms a synapse with the first dendrite also forms one with the second, is equal to 1*/c*. The uncorrelated case corresponds to *c* =*∞*, and *c* = 1 is the other extreme where all dendrites share the same set of input axons.

### Learning Rules

An algorithm similar to the passive-aggressive learning [30] is used to update the synaptic weights. There are two steps in the learning rule: first, dendrites are selected whose weights should be updated; second, the weights of the selected dendrites are modified so that the local activation reaches a target level *u*_target_. Specifically, the update on the *r*-th dendrite is given by

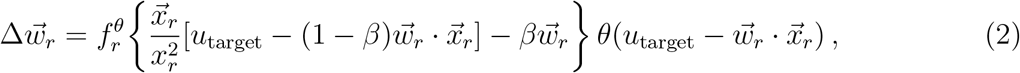

where *f*^*θ*^ is a binary mask function that decides which dendrites are selected, *β* is a decay factor that controls the L2 norm *w*_*r*_ of weight vector, and *θ* is a Heaviside function that prevents weight changes if the local activation already exceeds *u*_target_ before learning. The parameter *β* is chosen so that the distribution of *w*_*r*_ is centered around the target norm *w* = 1. After learning, dendrites that undergo weight changes have activations

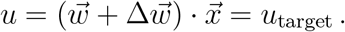

An intuitive picture for this learning rule is a diffusion process on the surface of an *N*_*s*_- dimensional unit hypersphere. The random walk step size scales as 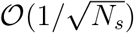. Whenever a new pattern is learned, the changes in weights interfere with the past memory. Reducing interference and achieving a high memory capacity requires a small random walk step size, which primarily depends on 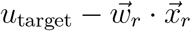 (terms with *β* only offer small corrections to 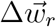 to constrain the weight vector around the unit sphere). This reveals two main factors that affect the capacity: *u*_target_ and the dendrite selection rule that determines the distribution of 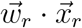 of the updated dendrites. There is a trade-off in choosing *u*_target_: it needs to be above the non-linear threshold *b* and large enough to achieve a strong response to learned patterns, but values that are too big lead to large learning step sizes that could severely disrupt past memories. Meanwhile, the selection rule should select dendrites that already have a fairly large activation for the new pattern in order to minimize the step size.

As a baseline rule, we consider a threshold-based selection rule (Fig. 1d left) with a constant

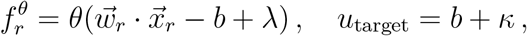

which means that only dendrites whose activations are above *b* − *λ* are selected, and the target level is always above the non-linear threshold *b* by a margin *κ*. This is a non-interacting rule because the update to each dendrite is independent of the activations of other dendrites. The number of selected dendrites, denoted by 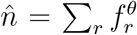, varies for different patterns. Given the target number *n* of selected dendrites, the value of *λ* is chosen so that 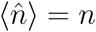.

There are two ways to introduce interactions among dendrites during learning: via dendrite selection or by adjusting *u*_target_. We consider one learning rule for each case. For the former, we still use *b* + *κ* for *u*_target_, same as in the baseline rule, but employ an n-winner-take-all (n-WTA, Fig. 1d right) selection rule — dendrites whose activations are among the top *n* are selected. For the latter, we set 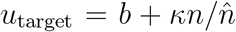, which depends on the number of selected dendrites, and we use the same threshold-based rule, except that if no dendrite is above *b−λ* (which might happen if *n* is small), then the dendrite with the largest activation is selected 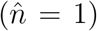.^1^ We call the former an n-WTA rule, and the latter a variable-*u*_target_ rule. In both rules the weight update to one dendrite implicitly depends on the activations of other dendrites. Before learning, a dendrite’s activation should rarely exceed the target *b* + *κ*, so we can omit the Heaviside function in eqn. (2). Then right after learning a pattern, the neuronal response to this pattern has expectation value ⟨*Y*⟩ = *nκ* for the baseline non-interacting rule, but is equal to *Y* = *nκ* for the interacting rules. This illustrates a crucial difference: interactions can regularize the neuronal response and result in a much smaller variance.

In biological neurons, both strategies for inducing dendritic interactions may be present, and they are likely implemented in a more complex manner than the simplified rules considered here. For instance, *u*_target_ may vary across both input patterns and dendrites, and dendrite selection may rely on mechanisms more intricate than the n-WTA rule (or threshold-based selection). Our aim, however, is not to capture biophysical mechanisms in full detail. Instead, we employ the rules introduced here as an idealized framework for isolating and analyzing the effects of dendritic interactions during learning.

Since the evolution of 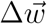 can be described as a random walk in an *N*_*s*_-dimensional space, the dynamics of the local activation *u* in response to a fixed input pattern *x*_0_ is a 1-dimensional random walk. For large *N*_*s*_, the evolution of *u* can be analyzed using the Fokker-Planck equation, which depends on the first few moments of Δ*u*. Immediately after learning an input pattern, the response distribution of *u* sharply peaks around *u*_target_. As other new patterns are learned, the center of the distribution gradually decays, and the spread becomes wider. The time evolution of this probability distribution is captured by its first two moments, *m*_1_(*t*) and *m*_2_(*t*). When inputs are sampled from the unit Gaussian distribution, these moments can be calculated as (Appendix A)

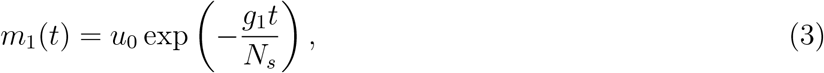

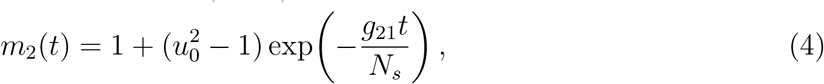

where *u*_0_ is the dendrite’s response to pattern *x*_0_ at *t* = 0, and the coefficients *g*_1_ and *g*_21_ are obtained from numerically solving a self-consistency equation, so that the distribution of the L2 norm of 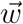 is centered around *w* = 1. The coefficients *g*_1_ and *g*_21_ characterize the response of an individual dendrite. These coefficients are *O*(*n/N*_*d*_), and their exact forms depend on the dendritic selection rule and the choice of hyperparameters (Appendix A). Therefore, the time constant of the random walk scales linearly with *N*_tot_ = *N*_*s*_*N*_*d*_. However, because dendritic signals are integrated at the soma to produce the output, the neuron’s response and its memory capacity exhibit a nontrivial dependence on *n*, and the dendritic non-linearity further complicates this dependence. In summary, the memory capacity scales linearly with *N*_tot_, and an appropriate choice of hyperparameters together with the dendrite selection rule will maximize the proportionality constant.

## Results

### Interactions increase capacity by reducing initial response variance

To compare the learning rules, we first consider a specific case where there are *N*_*d*_ = 300 dendrites, *N*_*s*_ = 200 synapses per dendrite, and the target number of selected dendrites to learn each pattern is *n* = 10. Unless otherwise stated, the same set of hyperparameters (*b* and *κ*) are used here, and they are optimized via a grid search for the baseline non-interacting learning rule. *β* is calculated separately in each rule to make the weight length distribution centered at *w* = 1 (Appendix A, and Fig. 2a). The distributions of the activations of selected dendrites before the weight update (Fig. 2b) are almost identical for the non-interacting and variable-*u*_target_ rules, and the first and second moments of the distribution for the n-WTA rule only differ by around 1%. The update step size Δ*w* has a similar mean for all rules, but the variable-*u*_target_ rule has a much larger variance (Fig. 2c). This agrees with our intuitive interpretation that the step size is approximately proportional to 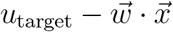, and since the statistics of 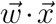 are similar, a large variance in *u*_target_ results in a high variance in the step size. Consequently, the weight norm distribution of the variable-*u*_target_ rule has a larger variance compared to the other rules (Fig. 2a).

**Figure 2.**
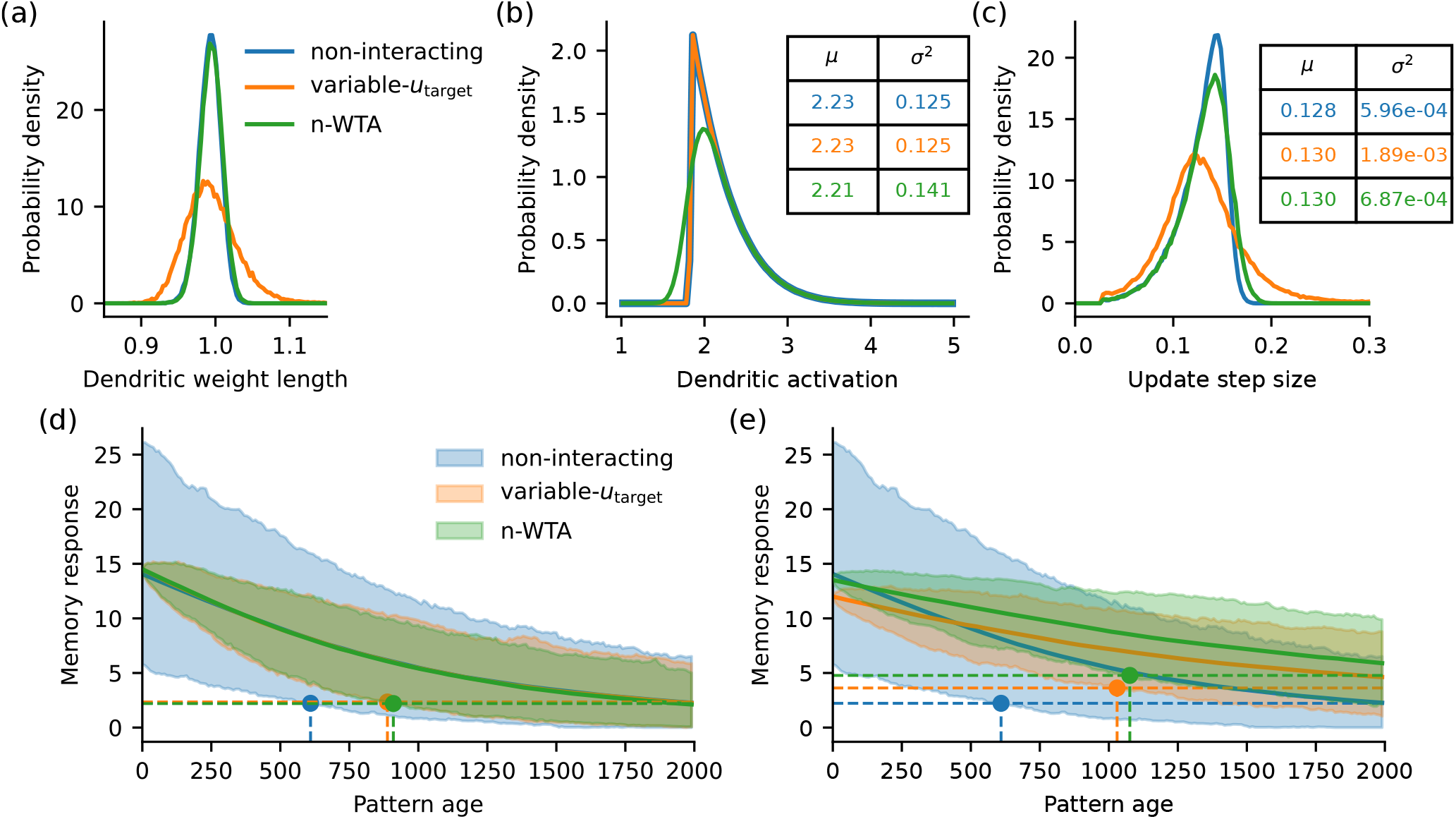
Comparison of learning rules (non-interacting: blue; variable-*u*_target_: orange; n-WTA: green) with uncorrelated inputs. In (a-d) all rules share hyperparameters optimized for the non-interacting rule, favoring it in the comparison. (a) Distributions of weight lengths. (b) Distributions of selected dendrites’ activation levels before learning. Inset: mean and variance of the distribution. (c) Distributions of weight update sizes. Inset: mean and variance of the distribution. (d) Temporal evolution of the memory response as a function of pattern age. Solid lines and shading indicate mean and 1%-99% interval (obtained from 1000 patterns). Horizontal dashed lines indicate the 99th percentile of the steady state distribution, vertical dashed lines indicate the memory capacity. (e) Same as (d), but with each learning rule using its own optimized hyperparameters.

Fig. 2d illustrates how the memory response distribution to a familiar pattern evolves with pattern age. We note that the curves for the mean response evolution overlap for all rules, which is consistent with the results that the mean update step sizes are similar. However, because the interacting rules have a much smaller initial variance of the response, they result in a much higher memory capacity compared to the baseline non-interacting rule. This difference can be further increased if hyperparameters are optimized separately for each learning rule (Fig. 2e).

### Interactions increase capacity by limiting weight correlation

So far we have compared learning rules using uncorrelated inputs, which corresponds to a shared-input coefficient *c* =*∞*. Next, we study the effects of interactions using inputs that are correlated across dendrites. We use the same parameter setting as in the previous section, which is optimized for the baseline non-interacting rule. Fig. 3a shows the evolution of response distributions when *c* = 3. The capacities of the different learning rules are all lowered compared to the case with uncorrelated inputs shown in Fig. 2d. However, the non-interacting rule is the most severely affected, with a 59% decrease in capacity, while capacities from the variable- *u*_target_ and n-WTA rule only decrease by 14% and 5%, respectively. The mean curves of the different distributions still overlap, so the main explanatory factor for the different capacities is again the variance of the response. Fig. 3b further shows how the capacities vary with the coefficient *c* that determines the input correlation. Notably, the capacity of the non-interacting rule eventually goes to 0 as the input correlation increases (smaller *c*), while the interacting rules still maintain high capacities even in the extreme case where *c* = 1.

**Figure 3.**
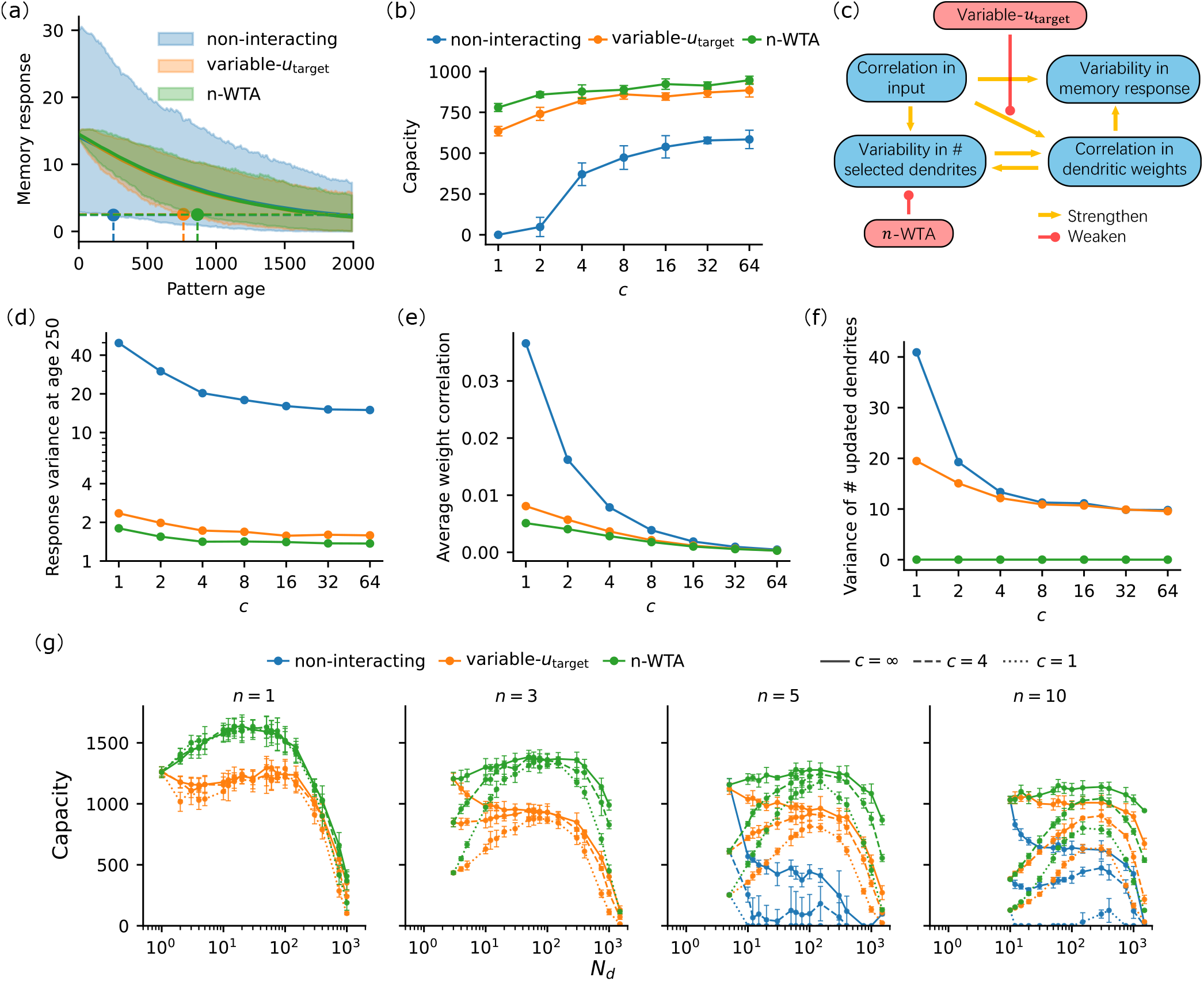
Comparison of learning rules (non-interacting: blue; variable-*u*_target_: orange; n-WTA: green) with inputs correlated across dendrites. In (a-b, d-f) all rules share hyperparameters optimized for the non-interacting rule, favoring it in the comparison. (a) Temporal evolution of the memory response as a function of pattern age (same format as Fig. 2d). (b) Memory capacities as a function of shared-input coefficient *c*. (c) Schematic showing how input correlations decrease capacity and how dendritic interactions can partially mitigate this reduction.(d) Variance of memory response (when pattern age is 250) as a function of shared-input coefficient. (e) Weight correlation as a function of shared-input coefficient. Correlation is computed in the input space (*N*_*a*_-dimensional). (f) Variance of the number of selected dendrites as a function of shared-input coefficient. (g) Memory capacity as a function of number of dendrites *N*_*d*_. From left to right, the target number of selected dendrites are 1, 3, 5, 10. Different linestyles correspond to different shared-input coefficients. The hyperparameters of each data point are optimized via a grid search. Error bars indicate *±*1 standard deviation across 6 trials.

Fig. 3c schematically explains how correlation in inputs lowers the capacity and how interactions alleviate this effect. As discussed above, the main cause of low capacity is the high variance in the neuronal response (Fig. 3d), which is directly caused by correlations in the inputs and correlations across dendritic weights (Fig. 3e). Since the input statistics are the same for the different learning rules, the difference in performance comes mainly from different levels of weight correlation. Note that we assume that there is no correlation across (axonal input) patterns; a pair of dendrites become systematically correlated in their weights only when they are driven by correlated inputs (for a given pattern) that arise from axons contacting both dendrites. Therefore, the weight correlation should approximately scale with the probability that a pair of dendrites is selected together for updating, *P*_co-select_. In an approximation where the dendrite selection is completely random, its expectation value is

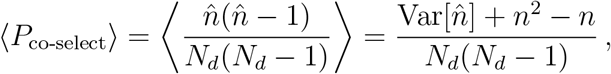

which increases with the variance of the number of selected dendrites (Fig. 3f), even though the average number is the same for all learning rules. In turn, for the threshold-based selection rule, correlation in inputs and correlated weights increase the variance of the number of selected dendrites — as the *u*_*r*_ values become correlated, they are more likely to stay above or below the selection threshold together. The n-WTA rule avoids this effect by directly constraining the variability in the number of selected dendrites. The variable-*u*_target_ rule, in contrast, can limit the correlation in weights by decreasing the strength of the weight updates when too many dendrites are selected.^2^ More intuitively, how much weights are driven by correlated inputs is reflected in the neuron’s output immediately after learning. Thus, regularizing the output interactions among dendrites limits the dendritic weight correlation, which results in a higher memory capacity.

So far, we have only considered a specific set of parameters that are optimized for the baseline rule in the uncorrelated input scenario. Next, we compare the learning rules for different values of *N*_*d*_, *n*, and *c*, while keeping the total number of synapses *N*_tot_ fixed, and the hyperparameters *b, κ* are optimized via grid search for each data point (Fig. 3g). For the non-interacting rule, the capacity drastically decreases as *n* decreases, and it reaches 0 when *n <* 5. For *n*≤4 the probability that 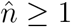 is less than 99%, and the false negative rate would always be above 1%, which according to our definition leads to a vanishing capacity. For both of the interacting rules the maximum capacity is achieved at *n* = 1, and the capacity curves are barely affected by the input correlation in this case. However, a biological neuron might not strictly select one dendrite to learn each pattern, since there are additional factors, such as noise, input correlations within a dendrite, etc. to consider, which could reduce the robustness of memories stored with *n* = 1. When the neuron uses one of the interacting rules, the capacity stays high for *N*_*d*_ values over a broad range, from tens to hundreds, and the capacity curves show clearer peaks in this range as the correlation in the inputs increases. Note that even under the most correlated inputs, the capacity can still remain at a high level for a wide range of *N*_*d*_ values. For *n >* 1, the capacity curves for different *c* values no longer overlap as tightly, but capacities around the peaks are still relatively close to each other, especially for *n* ≤ 5. In summary, when the neuron uses an interacting learning rule, high memory capacity can be achieved for a wide range of numbers of dendrites *N*_*d*_, input correlation levels, and *n* values.

### Robustness of results against model variations

The results presented in the previous sections were derived from a highly simplified parallel dendrite model. Although this model captures the crucial feature that dendrites act as basic computational units capable of learning and inference, it abstracts away many biological details. To assess the robustness and generality of our findings with respect to specific modeling assumptions, we performed similar simulations using several alternative model variants.

As a comparison baseline Fig. 4a presents the capacity curves under uncorrelated inputs using the original model and the n-WTA rule. (We only present results with the n-WTA rule here since it gives the highest capacity). In Fig. 4b we use the Heaviside step function instead of ReLU for the dendritic non-linearity. In Fig. 4c we use a sparse binomial distribution instead of a unit Gaussian distribution to generate inputs. Specifically, for each synapse on a dendrite the probability of receiving a non-zero input is 0.15 (sparsity), and the non-zero inputs are sampled from the binomial distribution Binomial 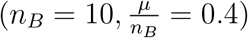, where *µ* can be interpreted as the average number of spikes in a burst and *n*_*B*_ as the maximum number of spikes in a burst.^3^ In Fig. 4d during recall we add input noise of the form *ση* independently to each synapse, where *σ* = 0.3 is the noise level and *η* is sampled from the unit Gaussian distribution. The capacity curves in Figs. 4b–4d exhibits qualitatively the same behavior as those from Fig. 4a. The shape of the curves and the relative ordering of curves for different *n* values are similar for these model variations.

**Figure 4.**
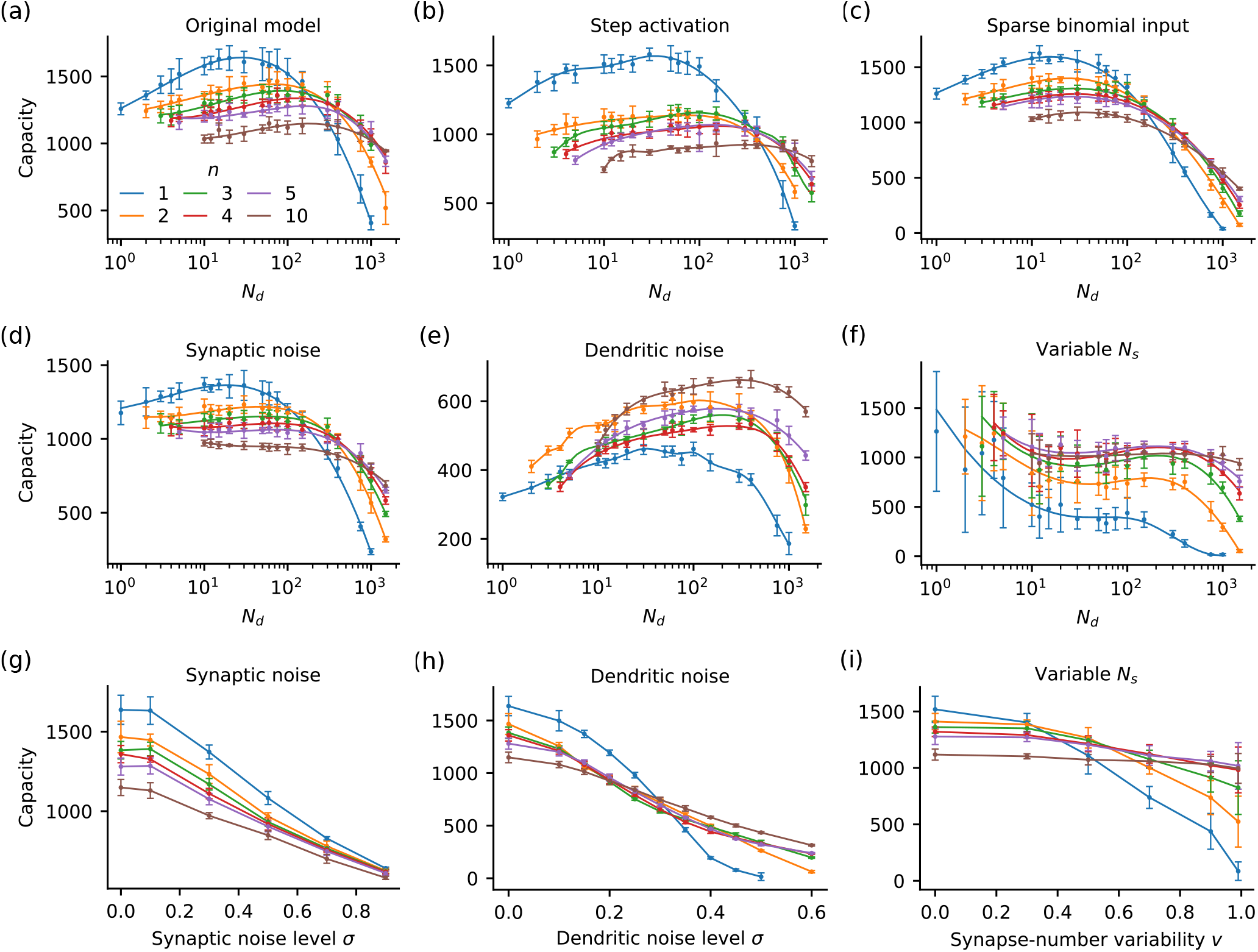
Robustness of memory capacity results to model variations. (a–c) Capacity vs. number of dendrites *N*_*d*_ under alternative baseline parameterizations/architectures (see the main text for details). The resulting curves exhibit similar qualitative behavior. (d–e) Capacity vs. *N*_*d*_ with synaptic noise (d) or dendritic noise (e). (f) Capacity vs. *N*_*d*_ when the number of synapses per dendrite is variable (heterogeneous dendritic sizes). (g–h) Maximum achievable capacity as a function of synaptic noise (g) or dendritic noise (h). (i) Capacity vs. the degree of synapse-number variability at fixed *N*_*d*_ = 100.

In addition to input noise, we also consider another type of noise, namely noise that is generated within the dendrite (Fig. 4e). It has the form *σκη*, where *σ* = 0.35 and *η* has the same definition as for input noise, but it is only added to dendrites whose activity *u*_*r*_ is above the activation threshold *b* (inside the dendritic nonlinearity) during recall. The noise is proportional to *κ* because a dendrite’s contribution to soma typically lies within [0, *κ*]. When dendritic noise is introduced into the model, the capacity curves (for different *n* values) not only decrease in magnitude but also change their relative ordering compared to the other model variants. The maximum capacity (across *N*_*d*_) for different values of *n* at various noise levels is shown in Fig. 4g for the input noise and Fig. 4h for the dendritic noise. The relative ordering of curves stays the same for a wide range of input noise levels, and *n* = 1 remains the highest curve. In contrast, for dendritic noise, the ordering is only the same for small noise levels, and larger *n* values actually lead to comparatively better capacity for increased noise magnitudes. This reflects two different strategies for a neuron to increase its signal-to-noise ratio (SNR) during memory recall in the presence of noise. For input noise, the noise approximately scales with 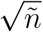, where 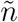 is the number of active dendrites during recall; for dendritic noise, the noise scales with 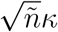. The signal approximately scales with 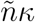. Therefore, in the presence of input noise, the SNR can be raised by increasing either *n* or *κ*, whereas for dendritic noise, the SNR can only be raised by increasing *n*. Since in a biological neuron there are various sources of noise, this result indicates that selecting *n* = 1 dendrite for learning is not always the best choice — the optimal choice depends on the noise level and noise type.

So far, we have only considered models in which all dendrites have the same number of synapses. In Figs. 4f and 4i, we instead introduce variability in the synapse number across dendrites: for each dendrite *r*, the number of synapses *N*_*s,r*_ is sampled uniformly from 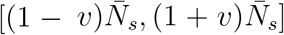, where *v* controls the degree of dendritic heterogeneity. We then define an effective dendritic length 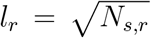, motivated by the fact that in our model the somatic signal is determined by fluctuations of the local activation, whose magnitude scales as 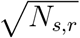. Accordingly, we make the dendritic weight norm and threshold scale with *l*_*r*_, and modify the learning rule so that the total update is distributed among the selected dendrites in proportion to their effective lengths (Appendix B). Similarly to the dendritic noise case (Fig. 4e), the relative ordering of the capacity curves for different *n* values in Fig. 4f differs from that of the original model, indicating that heterogeneity makes larger *n* more favorable. The noticeably larger variance at small *N*_*d*_ arises because, in this variant, only the average total synapse number is fixed, while the actually realized total synapse number fluctuates across the sampled neuronal model architectures, with 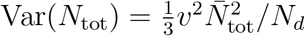(Appendix B).

Fig. 4i further shows that increasing the synapse number variability favors larger *n*, consistent with a failure mode when too few dendrites are selected: if the selected set of dendrite happens to contain only short dendrites, they must undergo disproportionately strong updates, which can severely disrupt previously stored memories. By contrast, for *n* = 10 the capacity is only weakly affected as *v* increases, suggesting that the modified learning rule can largely mitigate the effect of dendritic length variability when learning is distributed across sufficiently many dendrites. Together with the dendritic noise results, these results suggest that under more biologically realistic dendritic heterogeneity (and variability), memory performance can be improved by distributing learning across multiple dendrites, rather than relying on a single updated branch.

## Discussion

In this work, we compare three dendrite selection strategies for one-shot online familiarity detection in a parallel dendrite model neuron: a baseline non-interacting threshold rule with fixed *u*_target_, an interacting variable-*u*_target_ rule that rescales the update strength based on the number of selected dendrites, and an interacting n-WTA rule that fixes the number of updated dendrites per pattern. Although all three rules can be tuned to yield similar mean memory responses, the interacting rules achieve substantially higher memory capacity by limiting the variance of the neuronal response.

Biologically, the interacting rules studied here should be viewed as minimal abstractions of possible branch-level coordination mechanisms rather than mechanistic implementations. The n-WTA rule could approximate a competitive plasticity-selection mask in which strongly activated dendrites are preferentially recruited for synaptic modification, while weaker branches are excluded because of limited plasticity-related resources. The variable-*u*_target_ rule instead resembles a normalization of plasticity magnitude: when many dendrites are jointly recruited, a cell-wide or dendrite-population-level signal reflecting total postsynaptic activity could reduce the effective update per branch, whereas sparse recruitment would allow stronger modification of each selected branch. In this interpretation, the two interacting rules represent hard and soft forms of branch-level competition, respectively, both capturing the idea that plasticity at one dendrite need not be independent of plasticity at the others.

The key advantage of interactions is that they regularize learning at the level of the whole neuron, simultaneously limiting response variability and suppressing dendrite–dendrite weight correlations. Under the non-interacting rule, independent thresholding produces large fluctuations in the number of updated dendrites, which leads to a large variance in response. This effect becomes especially damaging when inputs are correlated across dendrites: correlated activations increase co-selection of dendrite pairs, driving correlations among dendritic weights and raising output variance. The n-WTA rule directly fixes co-selection statistics by constraining the number of updated branches, while the variable-*u*_target_ rule indirectly limits correlated weight growth by weakening updates when too many dendrites are recruited. As a result, interacting rules maintain high capacity even in strongly correlated input regimes where the non-interacting rule’s performance collapses.

We further test the generality of our conclusions under several model variations. The capacity for the n-WTA interacting rule remains high when replacing the ReLU with a step-like dendritic non-linearity, when using sparse spike-like inputs instead of Gaussian patterns, and when introducing input noise during recall. These results suggest that the benefits of interacting rules do not depend on a specific choice of input distribution or dendritic non-linearity, but instead reflect a more general effect of regularizing how plasticity is distributed across dendrites. At the same time, our analyses indicate that the optimal number of updated dendrites can depend on the sources of noise and heterogeneity: dendritic noise can favor recruiting multiple dendrites to improve signal-to-noise ratio, rather than selecting a single winning dendrite to update, and synapse-number variability can similarly shift the optimum toward updating multiple dendrites, since distributed learning is more robust to branch-to-branch differences.

The present work also has several limitations. First, we only model input correlation across dendrites via shared upstream axons, and do not consider correlation within a dendrite. Optimally incorporating within-branch correlations would likely require a further modified learning rule that accounts for local redundancy and competition for synaptic plasticity inside a branch. Second, the parallel-dendrite abstraction ignores the tree-like morphology of real dendrites, including branch points, signal attenuation, and location-dependent nonlinearities, all of which could shape the effective competition and plasticity allocation among dendritic branches. Testing whether the same interaction principles improve memory capacity in morphologically realistic dendritic trees is therefore an important next step toward identifying which forms of dendritic competition are plausible and effective in biological neurons.

## Appendices A Dynamics of 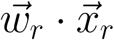

In this section, we will analytically derive the dynamics of local activation 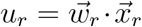 assuming that each element of the input is independently sampled from *N* (0, 1). We start from the update rule (2), copied here for convenience:

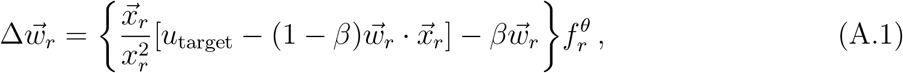

where the Heaviside function has been absorbed into 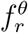. The parameter *β* (to be solved for self-consistently) is chosen so that the steady state distribution of the weight norm *w* is centered at *w* = 1. Since 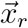 is drawn from a standard normal distribution, both the local activation *u* and its target value are *O*(1), and consequently, the terms within the square bracket in (A.1) are also *O*(1). Then, the magnitude of the update is 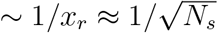, which means the width of the distribution of *w* will be at most 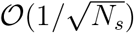 at steady state. Furthermore, at steady state, the weight update is approximately orthogonal to the weight vector, i.e.,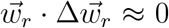, which implies that 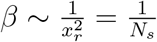.

Define 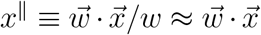, and define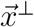 as the part of 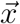 that is perpendicular to 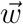, then the mask function is given by

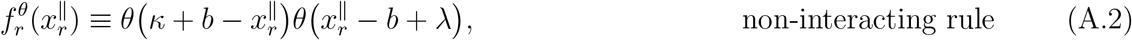

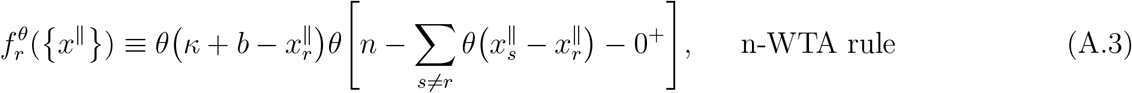

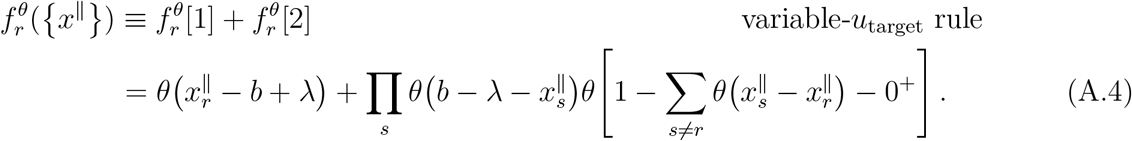

Here we omit the upper threshold for the variable-*u*_target_ rule for simplicity. The impact is minimal because *u*_*r*_ almost never exceeds *u*_target_ before learning. Then (A.1) can be rewritten as

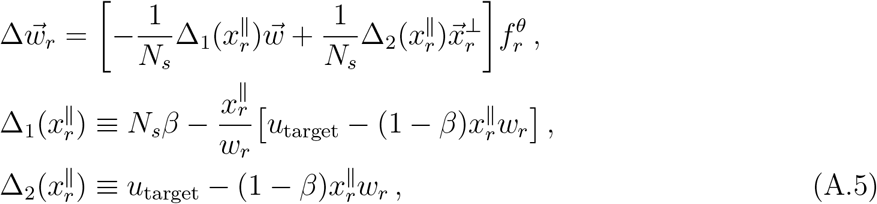

where *N*_*s*_ is used to approximate 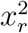. Since *w*_*r*_ *≈* 1 and 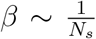, both Δ_1_ and Δ_2_ are *O*(1).

To apply the Fokker-Planck equation on dendrite *r*, we need to calculate moments of 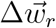 by integrating over the inputs 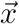. Note that now the calculation involves integrals over 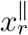 and 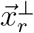, which can be done separately. To make the calculation simpler, we introduce a coordinate system 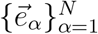 such that 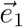 is parallel to 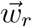 and all other basis vectors are orthogonal to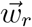.

Let the projection of 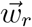 onto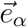 be denoted by 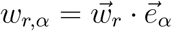. Then we have

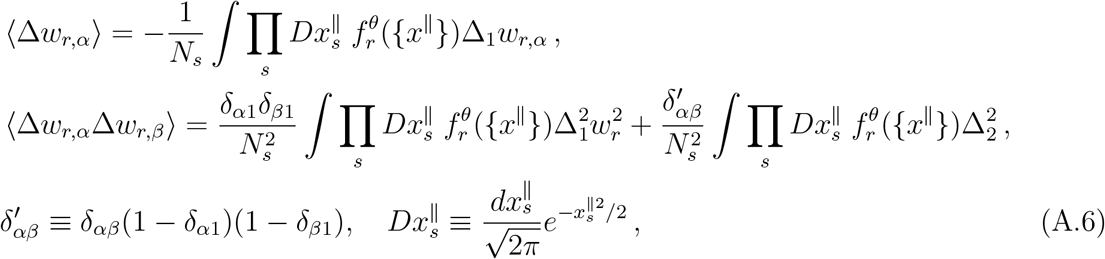

where we have used that integrals with odd powers of 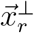should vanish. Since we only study the dynamics of the weights of a single dendrite, we do not consider cross terms like *⟨*Δ*w*_*r,α*_Δ*w*_*s,β*_*⟩* for *r≠ s*. Let 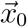 be the input to dendrite *r* at time 0. We are interested in the evolution of 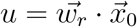. Moments of Δ*u* using moments of 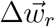 above can be expressed as

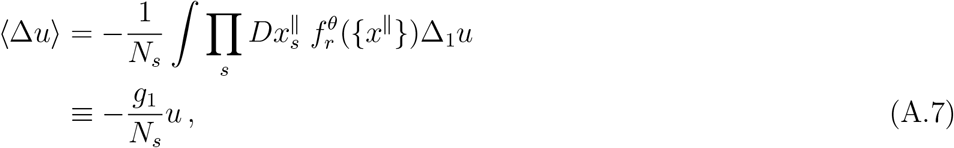

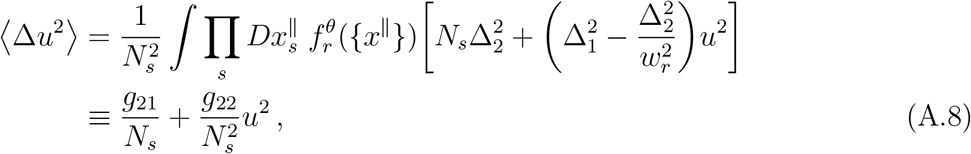

where we have used 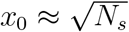. Remembering that Δ_1_, Δ_2_ are *O*(1), and that

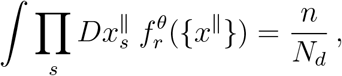

we can derive that *g*_1_, *g*_21_, and *g*_22_ are all *O*(*n/N*_*d*_). To calculate these coefficients more precisely, we can further reduce the high-dimensional integrals to 1-dim. or 2-dim. integrals that only involve 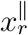 by computing all the other integrals.

For the non-interacting rule:

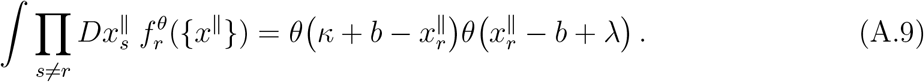

For the n-WTA rule:

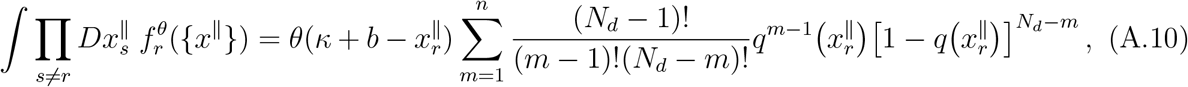

where *q*(*x*^∥^) is defined as

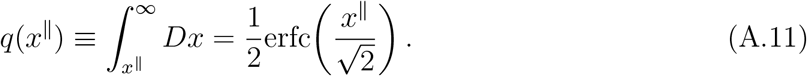

For the variable-*u*_target_ rule, reducing the integrals is more complicated because 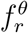 contains two terms, and the 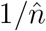 factor, which comes from *u*_target_ in Δ_1_ and Δ_2_, depends on all of the 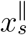. Consider 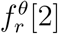 first. It corresponds to the simple case where all activations are below *b – λ* and 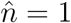:

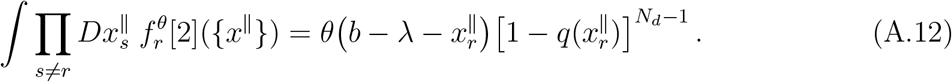

For the 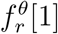 term, in (A.7) and (A.8) we will encounter terms with factors of 1, 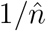, and 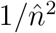. We need to consider them separately:

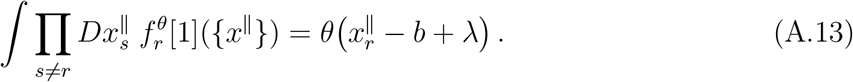

For the 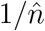 factor, noting that *q*(*b − λ*) = *n/N*_*d*_, we have:

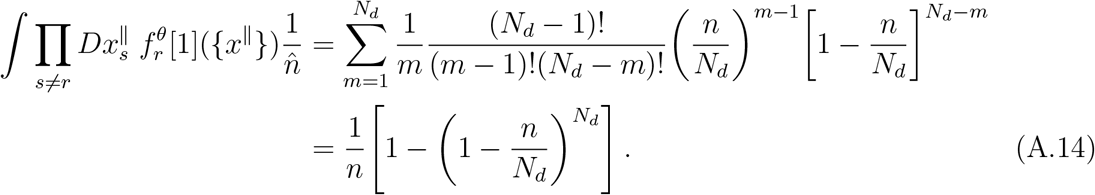

For the 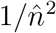 factor, by using identity

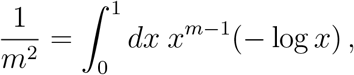

we have:

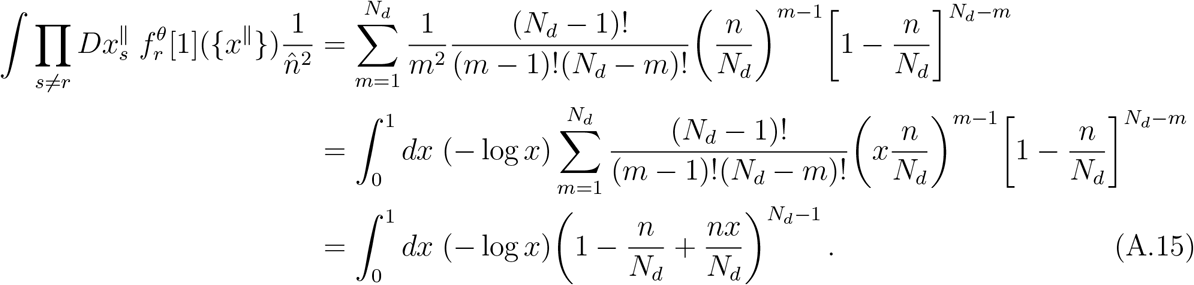

Therefore, using (A.9)–(A.15), the *g*_1_ calculation is reduced to a 1-dim. integral, the *g*_21_, *g*_22_ calculations are also reduced to 1-dim. integrals for the non-interacting and n-WTA rules, and they are reduced to 2-dim. integrals for the variable-*u*_target_ rule. All of these integrals can be numerically evaluated. Since we expect *w*_*r*_ to have a narrow distribution that centers at 1, it will be set to 1 for the numerical calculation.

The next step is to calculate the time-dependent distribution of *u*, which can be done by solving the Fokker-Planck equation:

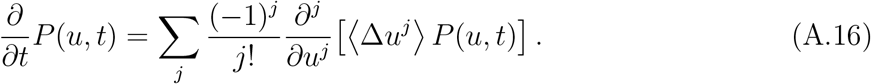

It is difficult to directly solve for *P*(*u, t*). Instead, we can use this equation to obtain the dynamical equations of moments of *u*. The *k*^*th*^ moment of *u* is

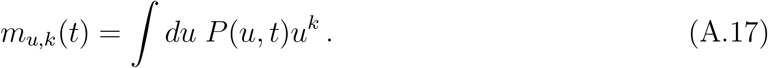

Plugging this into the Fokker-Planck equation gives

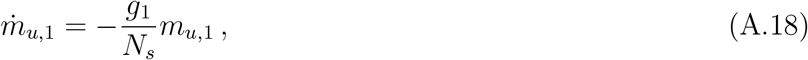

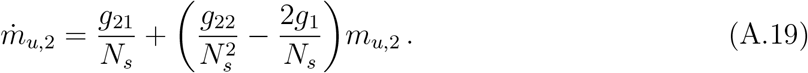

We can see that at steady state, *m*_*u*,1_ = 0, and *m*_*u*,2_ = *g*_21_*/*(2*g*_1_ *− g*_22_*/N*_*s*_). On the other hand, because *w*_*r*_ = 1 and 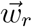 becomes completely uncorrelated with 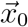 at steady state, we expect that 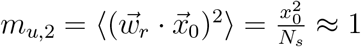 at steady state. Thus, we have

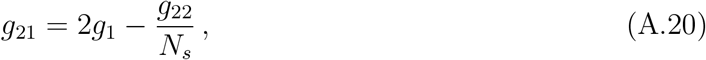

from which we can solve for *β*, the parameter that controls the length of 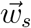_*s*_. Assume that at time 0, *u* is fixed at *u* = *u*_0_. With this initial condition, we can solve the differential equations above:

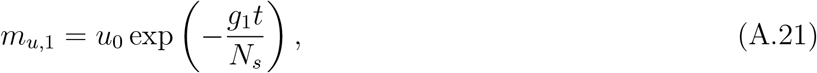

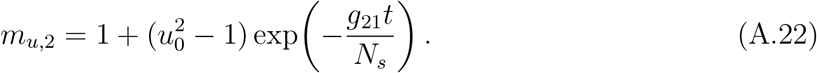

These equations characterize the evolution of dendrite’s response to a pattern starting from a given value: the mean response decays exponentially toward zero, while the spread of the response distribution approaches its steady-state value. Since *g*_1_ and *g*_2_ are of order *O*(*n/N*_*d*_), the characteristic time scale of this evolution is *O*(*N*_*s*_*N*_*d*_*/n*) = *O*(*N*_tot_*/n*). Thus, the overall memory lifetime scales linearly with *N*_tot_.

## B Generalization to variable dendritic lengths

In this section, we describe the modified model and learning rule used to incorporate variability in the number of synapses per dendrite. For each dendrite *r*, after sampling *N*_*s,r*_, define 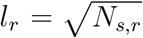, and 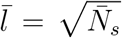, where 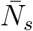 is the average number of synapses per dendrite. Then the neuronal response (1) is modified to

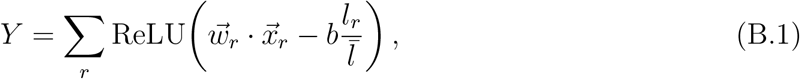

where the norm *w*_*r*_ should be approximately normalized to 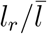. In other words, all activation related quantities are rescaled by a factor of 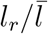. This choice is motivated by the fact that our model’s output is determined by the fluctuation of the local activation 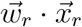, rather than by its mean (which can always be canceled by introducing a corresponding term in the dendritic threshold). If synapses on different dendrites have the same underlying strength distribution, then the fluctuation in *u*_*r*_ grows as 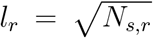. The constant 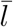 is introduced so that the generalized model is consistent with the original model at zero synapse-number variability.

Our guiding principle for this model variant is to preserve memory capacity by distributing the memory load approximately uniformly across synapses, so that longer dendrites carry a proportionally larger share of the memory load. This can in principle be achieved in two ways: either longer dendrites are updated more frequently while all dendrites undergo updates of comparable magnitude, or all dendrites are updated with comparable frequency while longer dendrites receive proportionally larger updates. Since it is difficult to control the update frequency of individual dendrites directly, we adopt the second strategy. Accordingly, as a variant of the *n*-WTA learning rule, dendrites are selected according to 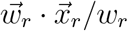, which has the same distribution for all dendrites. Note that any solution still need to satisfy the weight normalization constraint and that the neuronal response right after learning is *Y* = *nκ*. Therefore, a solution that properly distributes the weight update across dendrites can be obtained by solving:

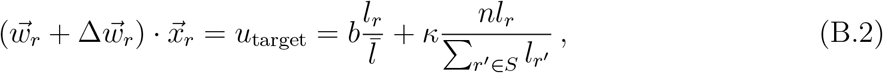

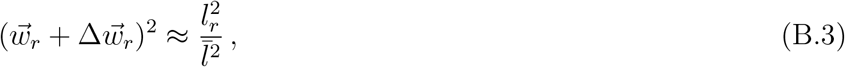

where *S* denotes the set of selected dendrites. To obtain an explicit update rule, we assume the post-update weight vector to lie in the span of the previous weight vector and the current input. Specifically, we first rescale the previous weight direction to the desired norm, 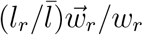, and then add the component along 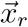 needed to satisfy (B.2) exactly. This gives

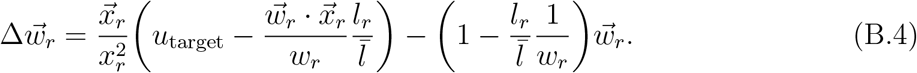

Substituting it into the expression for the norm gives

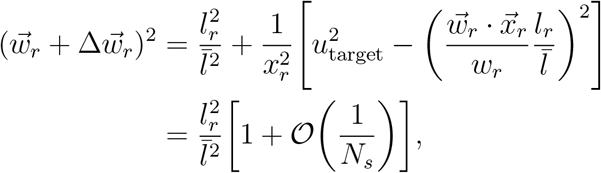

which indeed satisfies (B.3). Equation (B.4) is not the only possible solution: one could instead enforce (B.3) exactly, with or without an additional correction term, but the resulting expression is less transparent. With the present choice of update the deviation from the desired weight norm is an *O*(1*/N*_*s*_) correction.

A major difference of this learning rule compared to the original one is that there isn’t a constant decay factor *β* that determines the steady state weight norm distribution. Instead, without a decay process, (B.4) directly normalizes the weight to near 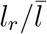 regardless of the initial weight length. To see the connection between these two learning rules, we can introduce scalars *λ*_1_, *λ*_2_ *>* 0, and relax (B.3) to

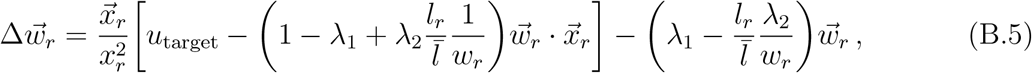

which still solves (B.2), but plugging it into (B.3) now gives

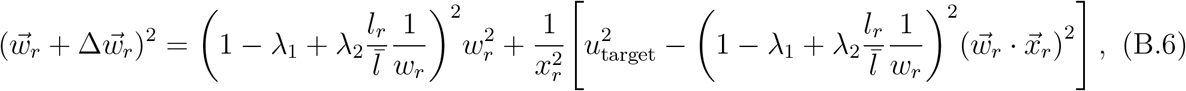

which leads to a relation for 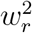. The original learning rule corresponds to *λ*_1_ = 1 *− β* and *λ*_2_ = 0.

In our simulations, *N*_*s,r*_ is sampled uniformly from 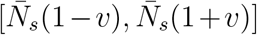. Thus, the variance of *N*_*s*_ is

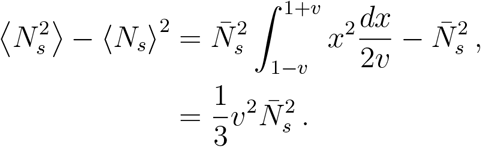

Therefore, the variance of the total number of synapses is 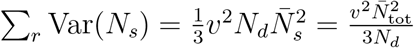.

## Footnotes

1 The 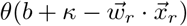 factor is omitted here for analytical convenience; the impact of this omission should be minimal because 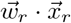 rarely exceeds *b* + *κ* before learning.

2 The weight correlation is driven by both the strength of the weight update and the probability that a pair of dendrites are co-selected. The weight update strength is approximately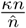. The pairwise weight correlation drive is thus approximately 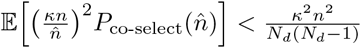, which is bounded by a value independent of the variance of the number of selected dendrites.

3 For this variant the learning rule also needs to be slightly modified: we use the centered (zero-mean) version of 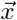 in (2), and impose a synaptic weight bound after learning each pattern to prevent extremely strong synapses.

